# Decoding real-world visual scenes from the human gamma band with flicker-evoked oscillations

**DOI:** 10.1101/2025.09.14.676118

**Authors:** James Dowsett, Inés Martín Muñoz, Paul Taylor

## Abstract

Current approaches to investigate the role of neural oscillations in natural scene processing have been limited to artificial stimuli and long data collection. We present a new way to decode real-world scenes participants are viewing from the steady-state visual evoked potentials (SSVEPs) evoked while wearing flickering LCD glasses. We discovered that SSVEP responses from real world scenes are surprisingly complex and have distinct waveform shapes: they differ markedly across scenes and participants but are consistent within individuals, even across multiple days. SSVEP shape varies greatly between stimuli, but is reliable, meaning that decoding works even with a single electrode. Decoding is highly accurate with 5-10 seconds of data and was still above chance level with less than a second of data. Decomposing the SSVEPs into frequency bands showed that the information about the visual scene is present across all of the harmonics of the flicker frequency, but with 40 Hz (gamma band) showing the highest amount of information across the different flicker frequencies tested. These findings implicate a broad range of oscillations in encoding real-world scenes, with a particular importance for 40 Hz. The SSVEP’s temporal profile is a rich source of information for decoding.

## Introduction

The ability to perceive complex 3D environments is a vital part of behaviour. Oscillatory neural activity is thought to support cognition by facilitating communication between distributed neuronal populations (Fries, 2005), but there is a currently a debate over which oscillatory frequencies - particularly, within the alpha and gamma bands – are most important for the perception of naturalistic scenes (Brunet et al., 2014; Chen et al., 2023; Hermes et al., 2015b). Yet the processing of complex real-world environments and situations may involve fundamentally different mechanisms to those elicited by small or static images on computer screens under lab conditions (Shamay-Tsoory & Mendelsohn, 2019; Snow & Culham, 2021). Cognitive neuroscience would benefit from studying neural activity in naturalistic (or semi-controlled) environments and exploring behaviour in real-world situations (Ladouce et al., 2017): however the limitations of current neuroimaging methods limit ecological validity in cognitive neuroscience (Gramann et al., 2014).

Recent work has begun to address this. For example, in a recent study (Dowsett et al., 2020) participants viewed real-world environments through LCD flicker glasses which allowed measuring the steady-state visual evoked potential (SSVEP). SSVEP phase and waveform shape varied according to how visual scenes was illuminated. Inspired by that, we here investigated whether the identity of real-world scenes can be decoded from SSVEP responses with LCD flicker glasses. With a waveform decoding approach, rather than focusing on specific features of the EEG signal (amplitude, latency etc), we test if the signal can be used to distinguish between the neural activity associated with different behavioural or cognitive states (in this case what the participant is looking at), and how much data is required to do so. An assumption-free, data-driven approach to brain time series analysis can reveal underlying brain states not reflected by, for example, power in canonical frequency bands (Balestrieri et al., 2025). If specific perturbations (e.g. flicker frequency) to a system (e.g. the brain) reveal more information encoded in that system (e.g. what the participant is perceiving), then the nature of these frequency specific perturbations can give some insight into the underlying mechanisms of how the system is encoding information.

Broadly, there are two ways to investigate the role of specific frequency bands of neural activity; firstly, we can filter the signal and ask which frequency band gives the most information about the mental state of interest (by far the most common approach), and secondly we can drive the brain at specific frequencies and ask which frequency results in the most information in the resulting EEG signal. In the current study we applied both approaches.

## Methods and Materials

In this pre-registered study, we set out to firstly test if it is possible to distinguish different real-world visual scenes with mobile EEG from SSVEPs generated with LCD glasses (figure 1). For the first experiment we used flicker at 10Hz, as the alpha band is known to evoke stronger SSVEP responses (Herrmann, 2001). In a second experiment we directly compared 10 Hz flicker with 1 Hz and 40 Hz, to look for evidence that there is some frequency specificity to the decoding accuracy.

**Figure 1:**
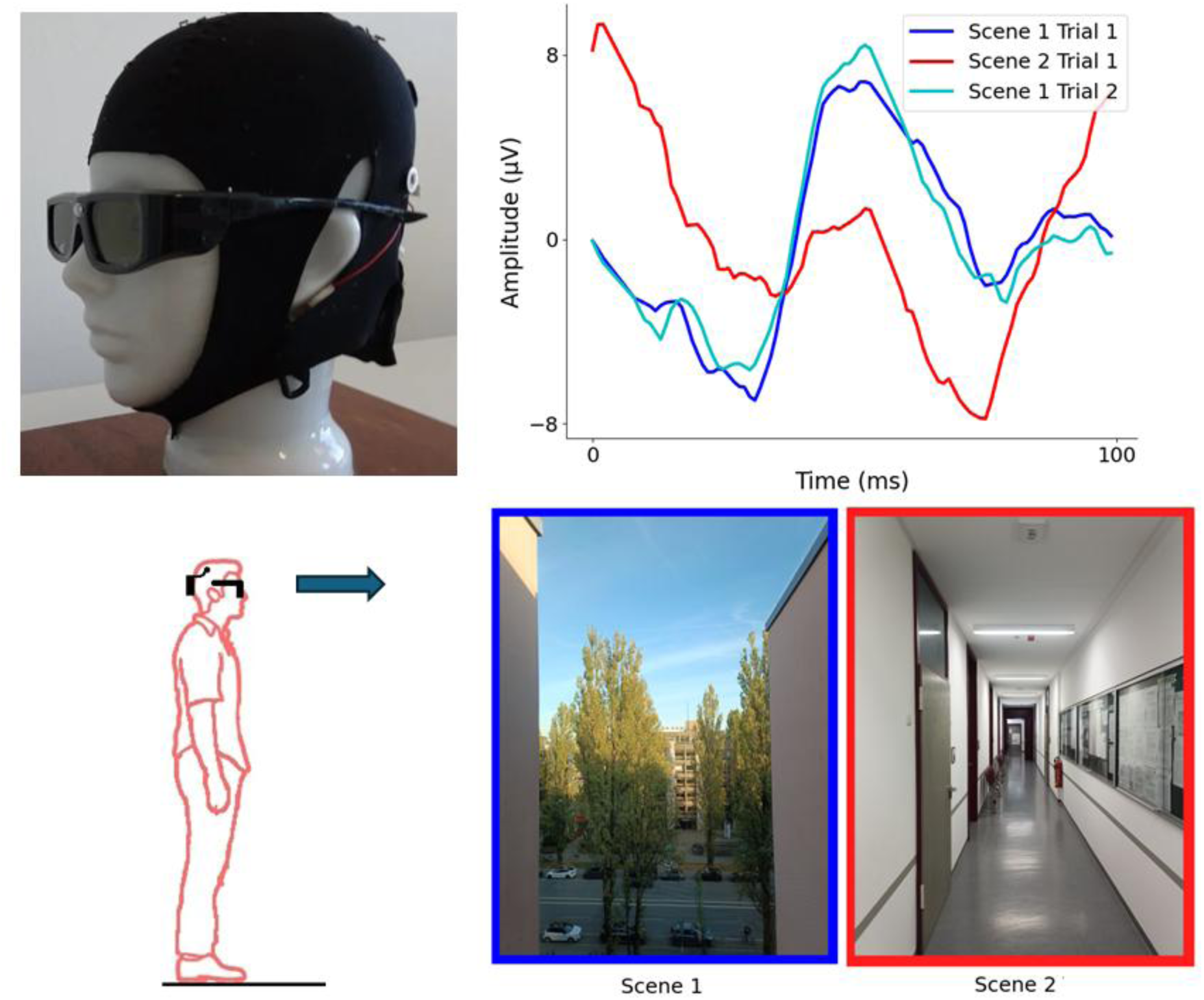
Top left; the LCD flicker glasses and EEG cap with amplifier. Bottom left; participants stood in various locations and fixated on a point in the distance whilst wearing the LCD flicker glasses and the mobile EEG. Bottom right; two example visual scenes used in the experiment from the perspective of the participant. Top right; Representative SSVEPs from one participant whilst looking at the two example scenes shown and a second SSVEP from scene 1 recorded at a later time. The correlation between the two SSVEPs from the same scene is high (0.93) and the correlation between the SSVEPs from different scenes is close to zero, allowing successful decoding.

### Participants

We recruited healthy participants between the ages of 18 and 30 years old. Participants were self-reported as right-handed, having no history of seizures or familial epilepsy, normal or corrected-to-normal vision and were not colourblind.

For Experiment 1, twenty participants were recruited (mean age: 24.85 years, STD: 2.9 years, min. 19 max. 30, 13 females). For Experiment 2, twenty new participants were recruited who had not participated in experiment 1 (mean age: 25 years, STD: 3.15 years, min: 19, max: 30, 13 females).

The local ethics committee (LMU Psychology, Munich, Germany) approved the study. All participants gave informed written consent before the experiment and were compensated 10€ per hour. The study was conducted according to the guidelines in the Declaration of Helsinki.

### EEG Acquisition

EEG data were recorded using an 8-channel mobile EEG system (LiveAmp, BrainProducts GmbH, Munich, Germany). The Ag-AgCl electrodes were in a cap and arranged according to the international 10-20 layout at O1, O2, P3, P4, P7 and P8 locations. Two EOG electrodes were used, one placed vertically below the right eye and another horizontally near the right eye’s outer canthus. EEG and EOG signals were digitised at 1000Hz, and the impedances were kept below 10kΩ. The reference electrode was placed at Cz, and the ground electrode at Fpz. The LiveAmp amplifier was attached to the back of the cap. The amplifier includes a 3-axis accelerometer which was used to detect head movements.

### Experiment 1

#### Procedure and Experimental Tasks

The objective of the first experiment was to determine whether SSVEPs can be used to discriminate between two different visual scenes. The EEG recordings were carried out at six locations in the Department of Psychology at the Ludwig-Maximilians University in Munich (figure 2). The locations were chosen intentionally to have distinctive characteristics. Two of the locations provided an outdoor visual field consisting of a natural scene observed at a far distance through a window on the fifth floor; the fixation point was a distance of approximately 75 m for one, and approximately 2.8 km in the other. The second two locations were indoors, with a medium-range visual field (fixation point approximately 10 m away) that covered a wide indoor space in one scene and a narrow hallway in the other. Finally, the last two locations provided a near-distance visual scene (approx. 1 metre) consisting of a flat surface; a blank white wall, and a wall with a painting depicting a stylised visual scene (i.e. a 2D surface with non-realistic depth cues).

**Figure 2:**
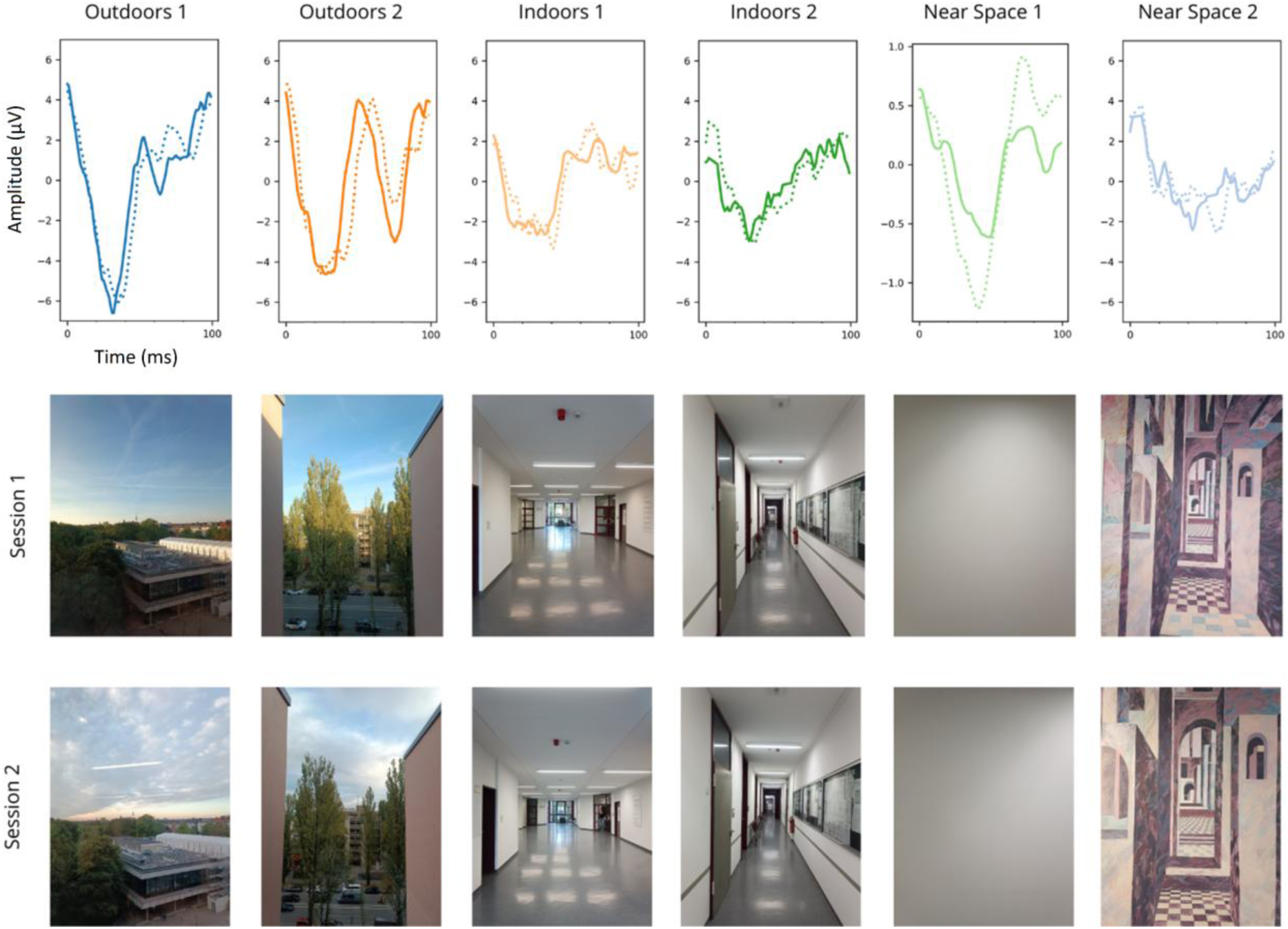
Example SSVEPs from one participant from each of the 6 locations used in experiment 1. Solid lines are the SSVEPs from session 1, dashed lines from session 2, recorded several days after session 1. Pictures show the visual scene from the perspective of the participant on both sessions. Differences in the visual scene, for example cloud cover and light source direction, can be seen across different recording days.

Each participant was required to complete two sessions, at least one day apart. Each session consisted of two 30-second trials per location. The data collected consisted of EEG and EOG recordings while standing at each location and receiving simultaneous flicker at 10Hz. Additionally, an eyes-closed seated recording of 30 seconds (without flicker) was performed in the laboratory to establish the individual alpha frequency of the participant. The order of the six different locations was pseudo-randomized for each trial.

For each location, the participants were instructed to stand at a specific point and to fixate on a designated point ahead of them. The participant initiated each trial by pressing a button attached to the flicker glasses.

Upon pressing the button, there was a baseline period of 5 seconds in which the participant looked at the fixation point in the visual scene, but no visual flicker was delivered. This was followed by 30 seconds of visual flicker. After 30 seconds of flicker stimulation there was another period of 5 seconds without flicker stimulation. A beep sound from a buzzer indicated the end of the trial.

#### Flicker stimulation

Flicker stimulation was delivered using custom-built liquid crystal display (LCD) glasses. An Arduino microcontroller (Arduino Nano) delivered voltage to the LCD glasses, turning the glass darker. The 10 Hz visual flicker was produced by changing the glass from transparent to semi-opaque using 5V pulses at 10 Hz with a 50% duty cycle (i.e. 50 ms dark, 50 ms transparent). A trigger aligned to each darkening of the glasses was sent to the EEG amplifier and used to segment the EEG and EOG data (Dowsett et al., 2020).

#### EEG processing

The data from each 30 second trial was segmented using the triggers from the flicker of the LCD glasses. This resulted in 300 segments, which were averaged to create a 100 ms SSVEP. For the main analysis, no filters or other pre-processing steps were applied. Subsequent analyses using filters are described below.

#### Visual Decoding

The waveform shape of each SSVEP was compared to the SSVEP from the same location and to each of the other locations, with a Pearson’s correlation (i.e. a correlation coefficient of close to 1 would indicate the waveform shapes are highly similar), here after referred to as correlation decoding (figure 1). We developed a score-based decoding algorithm to quantify whether different visual scenes can be decoded from their associated SSVEPs. For each location, a baseline correlation is calculated between the SSVEPs of the first and second trials at that specific location. This typically showed a high correlation, indicating a good test-retest reliability (median correlation from electrodes O1 and O2 across all locations = 0.91). This correlation was then compared to the correlation between the second trial at that location and the first trial at each other location individually; a point was awarded to that location if the baseline correlation was higher than the test correlation (i.e. successful decoding). This process was repeated for all locations, resulting in each electrode receiving a score out of 5 for each of the 6 locations (30 in total). A score of 5 out of 5 would indicate that the SSVEP can be successfully discriminated from all the other scenes due to it having a waveform shape which is both distinct from the other locations and consistent across trials.

To assess the statistical significance of the correlation decoding scores, we used randomly generated simulated data. For each comparison between baseline correlation and the test correlation, the outcome was randomly determined with a 50% chance of being correct for each location for each electrode and participant. This process was repeated 100,000 times to create a distribution of the score for the null hypothesis, with the average score being 50%. A Z-score and associated p-value were calculated for all possible scores between the maximum score and the chance level score. The alpha level (α=0.05) was corrected using Bonferroni correction for multiple comparisons for all 8 electrodes, giving a corrected alpha of 0.05/8 = 0.00625. This procedure indicated that an average decoding score above 55.1% across 20 participants is significant corresponding to the Bonferroni corrected p-value.

#### Comparison of decoding accuracy within one session to decoding accuracy across multiple days

For a neuroimaging methodology to be useful in real-world situations it needs to be robust to variability in the environment, in the participant, and in the exact recording conditions. We compared the correlation decoding from data collected on the same day (within session) to decoding across multiple days (between sessions), which would contain some incidental differences in the visual scene e.g. differences in light levels due to time of day, cloud cover etc. (figure 2). For this comparison, we took the average decoding accuracy across all EEG electrodes (O1, O2, P3, P4, P7 and P8) for all 6 visual scenes, to create one average decoding score per participant. The same was then done for the decoding across multiple days. A permutation test was performed with the following procedure: the difference between average decoding across participants between sessions was subtracted from average decoding within sessions. The labels (within or between sessions) for each participant were then randomly either switched or left unchanged, and the average was computed again, and this was repeated 100,000 times to create the null distribution. The *p*-value was calculated from the normalised distance of the actual value from the distribution of the random values (Cohen, 2017).

#### Effect of removing eye-blinks and movement artefacts

Typically, EEG analysis pipelines need to compensate for eye-blinks, eye-movement and head/body movement, to get a clean signal. A significant advantage of SSVEPs is the high signal-to-noise ratio, which we have previously demonstrated is sufficient to get a clean signal during walking (Dowsett et al., 2020). We expected that the decoding accuracy is highly resilient to occasional eye-blinks and small movements. To test this, we directly compared the decoding accuracy with and without removal of segments corrupted by eye-blinks and movements.

For the eye-blink/movement removal, 100ms segments of data were rejected from the averaging if the corresponding segment in the vertical EOG or horizontal EOG channel had a peak-to-peak amplitude greater than 100μV, which was determined by visual inspection to indicate a segment occurring during an eye-blink or eye-movement. Corresponding segments of data from the head-mounted accelerometer were checked in the same way and the EEG segment was rejected if the acceleration varied by more than 30 mg in any direction, indicating the participant was moving. The resulting decoding scores were then compared to the decoding scores without removal of artefacts.

#### Effect of 50 Hz notch filter on decoding accuracy

A possible confound in the decoding accuracy might be 50Hz line noise, which is a harmonic of 10Hz and would in theory sum in phase with the flicker. In theory the 50 Hz line noise could be higher in certain locations than others. However, this is highly unlikely to affect the decoding accuracy because the flicker would begin at an arbitrary phase in the 50Hz noise with each trial, and as such would not contain any useful information. As an additional check to confirm there was no effect of line noise, we compared the decoding accuracy before and after filtering the data using a 50 Hz notch filter (second-order IIR notch filter with a quality factor of 30).

#### Comparison of decoding with waveform shape vs. peak-to-peak amplitude

Studying SSVEPs using the waveform shape/phase is a fairly novel approach, whereas in the majority of SSVEP studies only the amplitude of the SSVEP is considered. We performed an equivalent decoding analysis using the peak-to-peak amplitude of the SSVEP and directly compared this to the correlation decoding of waveform shape.

For each location, we calculated the difference between the peak-to-peak amplitude of the SSVEP from trial 1 and trial 2. This amplitude difference was then compared to the peak-to-peak difference between the second trial at that location and the first trial at all other locations, one at a time. The scoring was the same as in the correlation analysis: a point was awarded to that location if the peak-to-peak amplitude difference was smaller than the amplitude difference with the SSVEP from the other location (successful decoding). This process was repeated for all locations, resulting in each electrode receiving a score out of 30 as above. A permutation test was performed to test for differences between correlation and amplitude decoding, following the same procedure as described above.

#### Determining the amount of flicker time needed to successfully decode

For the main experiment we recorded 30 s of data with 10 Hz flicker and attempted to decode the visual scene. Assuming this was successful, we then set out to ask how much data was necessary to decode the visual scene. For this analysis we ran the same correlation decoding procedure described above but used progressively less data. We used the full 30 seconds, followed by the first 25, 20, 15, 10, 5, 4, 3, 2, 1 seconds, and then sections of data less than 1 second removing 0.1 second each time, until the decoding accuracy fell to chance level. This was done for both the inter-trial decoding, and the inter-session (i.e. across different days). This was repeated for each electrode.

#### Bandpass filtering

The process of averaging segments the length of the period of the flicker is equivalent to a comb filter; the signal is a combination of the neural response at the flicker frequency and all the harmonics of that frequency. To further investigate the contribution of different frequency bands to the decoding accuracy we applied band-pass filters to the EEG data before making the SSVEPs and repeated the correlation decoding procedure described above for each. A 4th order Butterworth bandpass filter was applied at ±1 Hz around each harmonic, i.e. at 10, 20, 30, … up to 240 Hz using the SciPy Python library (version 1.10.0).

#### Comparison of decoding using endogenous alpha power without flicker

We performed further analysis on the 5 seconds of EEG data before the flicker started, but while the participant was fixating, to determine if the power of the endogenous alpha oscillations was significantly different from one scene to the next, and if this data could be used to decode the visual scene. For this analysis the resting data was segmented into a 4 second segment starting 1 second after the trial started (to avoid any movement artifact from the participant pushing the button to start the trial), and this was baseline corrected, multiplied by a Hanning window, and an FFT was applied. The maximum amplitude between 8 and 14 Hz was taken as the alpha peak for each trial (a 4 second segment gives a frequency resolution of 0.25 Hz). These peak values were then used for the decoding algorithm described above, where correct decoding would result from the location having a more similar peak alpha amplitude across trials than for any other locations.

#### Relationship between individual alpha frequency, flicker frequency and decoding accuracy

Previous studies have shown a relationship between the neural effect of visual flicker and how close this flicker frequency is to the participant’s individual alpha frequency (Nuttall et. al 2022, Jaeger et. al. 2023). We wanted to investigate any possible relationship between this frequency difference and decoding accuracy. The IAF of each participant was calculated for each session using the average FFT spectrum of electrodes O1, O2, P3, P4, P7 and P8. The IAF is determined as the frequency of the maximum peak in the alpha range, 8 to 14 Hz (frequency resolution of 0.25 Hz). Once the IAF had been calculated for each participant, the absolute difference between the IAF and the flicker frequency (10 Hz) was calculated. Additionally, the average visual decoding score of each participant for each session was calculated, excluding EOG and accelerometer channels. The decoding score was calculated and summed across all six locations, six electrodes and 5 comparisons per location, resulting in a maximum score of 240. The Pearson correlation between the visual decoding scores and the absolute difference between IAF and flicker frequency was calculated and tested for significance.

#### Relationship between decoding accuracy across days and differences in luminance

During each trial, the luminance level at the location was recorded on a smartphone with the “Lix Light Meter Pro” app (Google Play Store, Android). The app uses the light sensor on the smartphone to record the luminance measurement. The centre of the sensor was aligned with the fixation point and a luminance value was recorded for each location on each day of recording, at the time of data collection. For inter-session trials the luminance value on day one was subtracted from day 2. The Pearson correlation between the visual decoding scores and the difference in luminance was calculated. We hypothesised that a larger difference in luminance would result in a lower decoding accuracy, i.e. a negative correlation.

#### Decoding individual participants

The SSVEP waveforms were not only unique for each visual scene, but also unique across participants. For any one visual scene the SSVEPs from all participants were distinct. This is quite unlike ERP experiments where the evoked responses are sufficiently similar that they can be averaged across participants to produce a grand average. In the case of SSVEPs of real-world scenes, the SSVEPs are different phases and waveform shapes, meaning that any group average would be close to zero and would be meaningless. This individual variability allows us to not only decode the scene based on one participant’s previous SSVEPs, but also to distinguish which individual the data came from in trial 2, given the SSVEPs from all participants in trial 1. We ran a similar correlation decoding procedure to the one described above. The SSVEPs from all 20 participants for each location in trial 1 were compared to the SSVEPs from the same location in trial 2. For each location a point was awarded if the SSVEP for a participant from trial 2 had a higher correlation with the SSVEP from trial 1, than with the SSVEP from trial 1 for any of the other participants. For 20 participants the chance level would be 5%.

### Experiment 2

#### Flicker stimulation

The second experiment aimed to determine whether different visual flicker frequencies can also be used to decode the visual scene. We selected three of the six locations from experiment 1 (one far, one medium distance, and one near) and performed two trials at each location, with visual flicker stimulation at 1, 10 and 40Hz. An eyes-closed recording was also performed to establish the individual alpha frequency of the participants. In this experiment, the stimulation period was 10 seconds for the 10 Hz and 40 Hz stimulation and 100 seconds for the 1Hz stimulation. This was done to keep the number of flickers constant across the 1 Hz and 10 Hz conditions (i.e. in each case, the glasses flickered 100 times). A rest period of 5 seconds served as a baseline before and after the visual stimulation.

For the 1 Hz condition we kept the length of the dark period of the flicker the same as the 10 Hz condition: 50 ms dark followed by 950 ms transparent (i.e. a 5% duty cycle). This is because a 1 Hz flicker with a 50% duty cycle is perceptually very different from higher frequency flicker and would have resulted in separate evoked responses from the darkening and the glass going transparent 500 ms later. We wanted to keep the visual stimulation as similar as possible while only changing the flicker frequency.

For the 40 Hz condition we kept the stimulation duration constant (10 seconds), which resulted in the glasses flickering 400 times per condition. The duty cycle for the 40 Hz condition was 50 % (i.e. 12.5 ms dark and 12.5 ms transparent). At this frequency the flicker is barely noticeable, lying just below the standard flicker fusion frequency. The second experiment was conducted in a single session, and the order of the stimulation frequencies and the three locations was randomised across participants.

#### Comparison of Decoding accuracy across flicker frequencies

The correlation decoding accuracy was calculated using the same procedure as in experiment 1, except in this case there were only 3 locations (i.e. a maximum score out of 6, two for each location) and 3 different flicker frequencies. Because the results of experiment 1 showed fairly similar decoding accuracies from each of the EEG electrodes, we averaged the score across the 6 EEG electrodes, resulting in one score out of 6 for each flicker frequency. The Bonferroni corrected alpha level was 0.05/3 = 0.0166. We ran a simulation as in the first experiment to determine statistical significance; this time the simulated null data indicated that an average decoding score above 54.17% is significant corresponding to the Bonferroni corrected p-value. Note that this is a different value from experiment 1 because there were only 3 locations and we averaged across electrodes. To compare the decoding accuracy of the 3 flicker frequencies we used permutation tests (as described above) to compare 1 Hz vs 10 Hz, and 10 Hz vs 40 Hz.

#### Band pass filtering

As in experiment 1 we wanted to investigate the relative contribution of different frequency bands to the decoding accuracy. For the 10 Hz and 40 Hz conditions we performed the same procedure as in experiment 1: band-pass filters +/- 1 Hz around each of the harmonics of the flicker frequency were applied before averaging, and then the decoding was performed as before. For the 1 Hz condition the evoked potentials were complex waveforms which were not necessarily limited to the exact harmonics of the flicker frequency. This is because 1000 ms is enough time for various oscillations to be evoked by the 50 ms darkening of the glass which are not exact harmonics. For this reason, we investigated the relative contribution of different frequency bands in the 1 Hz SSVEPs with a series of wider band pass filters, each covering a 10 Hz band centred around 10, 20, 30, … etc. Hz.

### Results: Experiment 1

#### Visual Decoding

At the group level, and using the correlation-based decoding algorithm, we obtain median decoding accuracies above 90% for all EEG electrodes (figure 3). All decoding scores for EEG and EOG electrodes were considerably higher than the 55.1% accuracy needed to achieve significance at the Bonferroni corrected value (*p* = 0.00625). All electrodes resulted in Z scores of greater than 19.65 (corresponding to 90% median decoding accuracy), with all *p* values lower than 2.99 × 10⁻⁸⁶.

**Figure 3:**
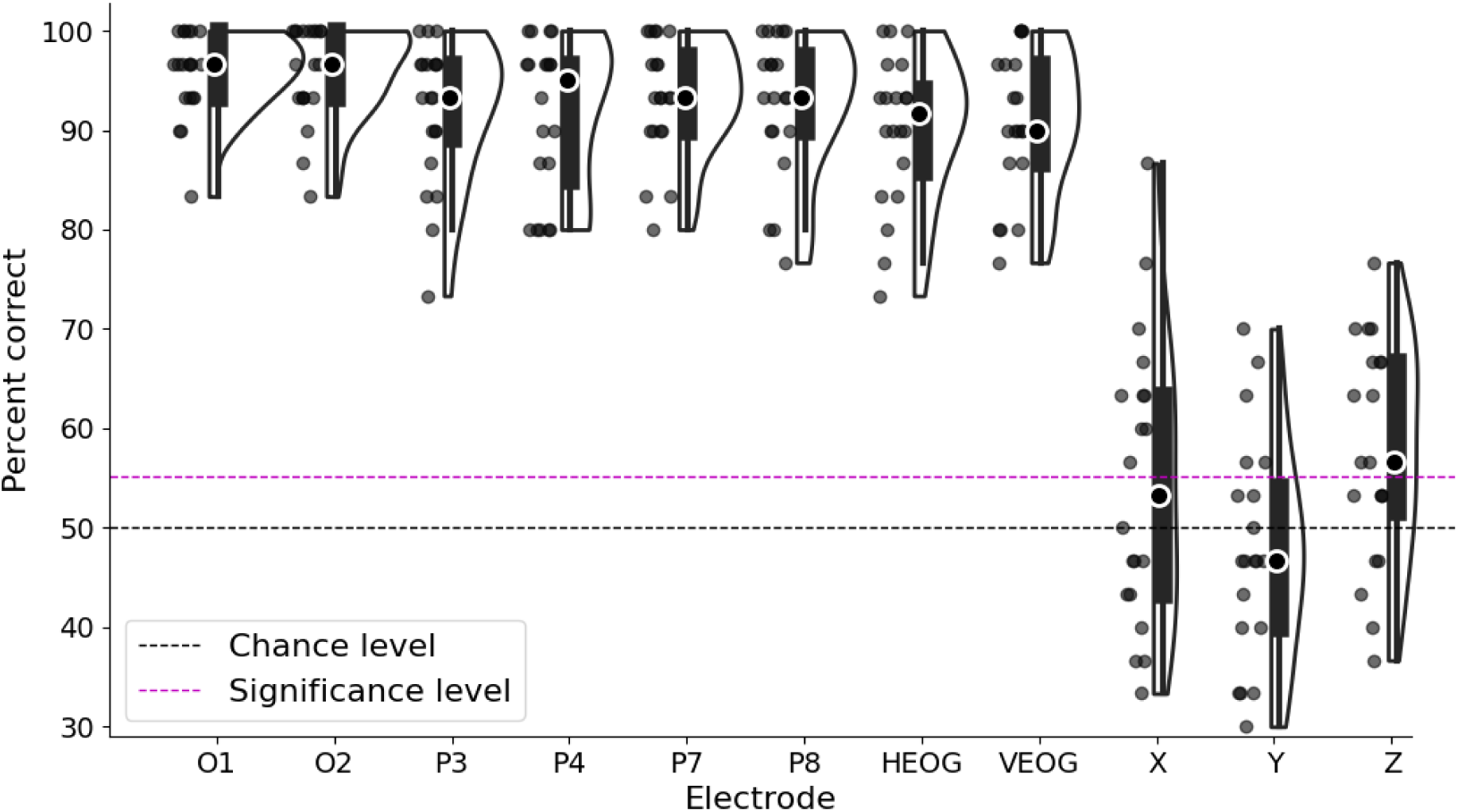
Decoding accuracy in percent of visual scenes correctly identified for all participants and all electrodes/sensors in experiment 1, within sessions. Each dot indicates one participant, black dot with white outline indicates median score, black dashed line indicates 50% chance level for pairwise decoding, red dashed line indicates the group average percentage correct corresponding to a Bonferroni corrected p-value of 0.00625. All EEG and EOG electrodes show well above chance level decoding accuracy. Evoked potentials from accelerometer sensors in the X, Y and Z directions are close to chance level decoding.

Additionally, we see that the accelerometer channels along all three directions have a decoding accuracy near chance level, confirming that the visual scenes cannot be decoded simply from the head movement of the participant time-locked to the visual flicker.

The electrodes with highest accuracy were occipital (O1 and O2), with 9 out of 20 participants scoring 100% for electrode O2, which indicates perfect decoding across all 6 scenes from this electrode.

#### Comparison of decoding accuracy within session to decoding accuracy across multiple days

When averaging across all EEG electrodes the permutation test revealed a significant difference between the median decoding accuracy within session (93.4%) and across multiple days (81.7%): Z-score, 3.30, *p*-value, 0.00048. The decoding accuracy across multiple days was still highly significant, with all electrodes having a median score of above 79% accuracy (figure 4), which corresponds to a Z score of 14.2 and a *p* value of 4.13 × 10⁻⁴⁶.

**Figure 4:**
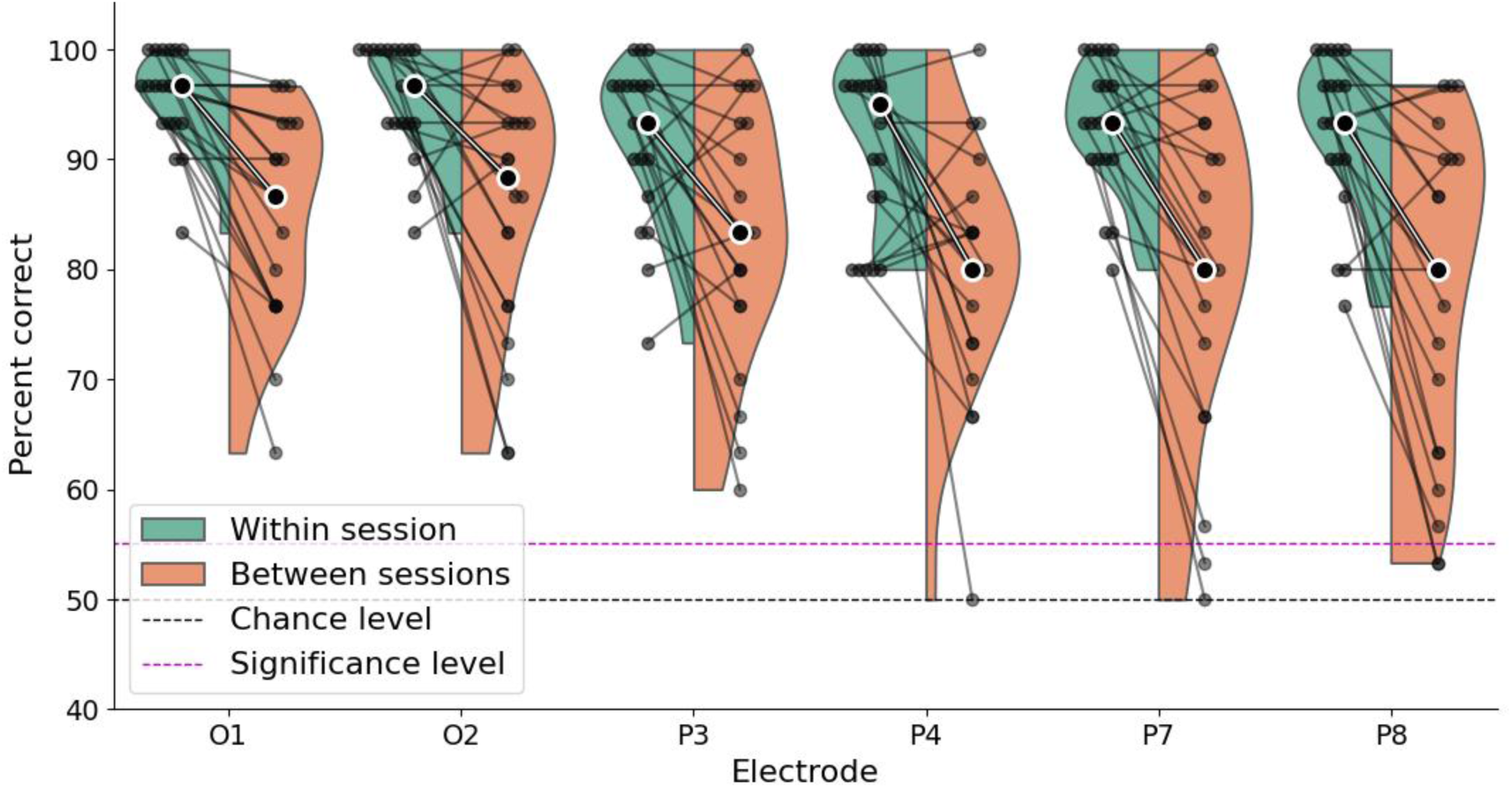
Decoding accuracy in percent of visual scenes correctly identified for all participants and all EEG electrodes in experiment 1 comparing within sessions (i.e. same day) and between sessions (i.e. across multiple days). Each dot/line indicates one participant, black dot with white outline indicates median score, black dashed line indicates 50% chance level for pairwise decoding, red dashed line indicates the group average percentage correct corresponding to a Bonferroni corrected p-value of 0.00625. All EEG and EOG electrodes show well above chance level decoding accuracy for both within and between sessions. Within sessions decoding is consistently better than between sessions.

#### Effect of removing eye-blinks and movement artefacts

The removal of segments of data containing eye-blinks, eye-movements and head movements made very little difference to the decoding accuracy. The mean difference in decoding accuracy across participants, trial and electrodes was 0.28% for within session decoding, and the maximum average difference for any one electrode was 0.67%. This is due to eye-blinks and movements being sufficiently rare/small that they did not modify the waveform shape of the SSVEP, because they account for a very small percentage of the overall number of segments. We therefore concluded that removal of eye blinks was not necessary for all subsequent analyses.

#### Effect of 50 Hz notch filter on decoding accuracy

The effect of the 50 Hz notch filter on the decoding accuracy was very small: The average difference in decoding accuracy across participants, trial and electrodes was 0.07% for within session decoding, and the maximum average difference for any one electrode was 0.5%. Based on this we can conclude that the decoding accuracy was not dependent on, or affected by, 50 Hz line noise. These small differences may have been due to neural activity in a 50 Hz harmonic of the SSVEP itself. The 50 Hz notch filter was deemed unnecessary and was not used in any further analyses.

#### Comparison of decoding with waveform shape vs. peak-to-peak amplitude

Decoding accuracy using the correlation of waveform shape was consistently better than decoding using the peak-to-peak amplitude of the SSVEPs (figure 5). For the within session results the average decoding accuracy across electrodes and participants was 92.69% with the correlation method and 70.48% with the peak-to-peak method. Between sessions the decoding accuracy was 81.5% with correlation decoding and 63.4% with amplitude decoding. The permutation tests showed a significant difference between correlation decoding and amplitude decoding for both within sessions (Z = 4.36, *p* = 0.000013) and for between sessions (Z = 4.05, *p* = 0.000052). This demonstrates that the waveform shape of an SSVEP (when using LCD glasses) contains significantly more information about the visual scene than the amplitude of the SSVEP.

**Figure 5:**
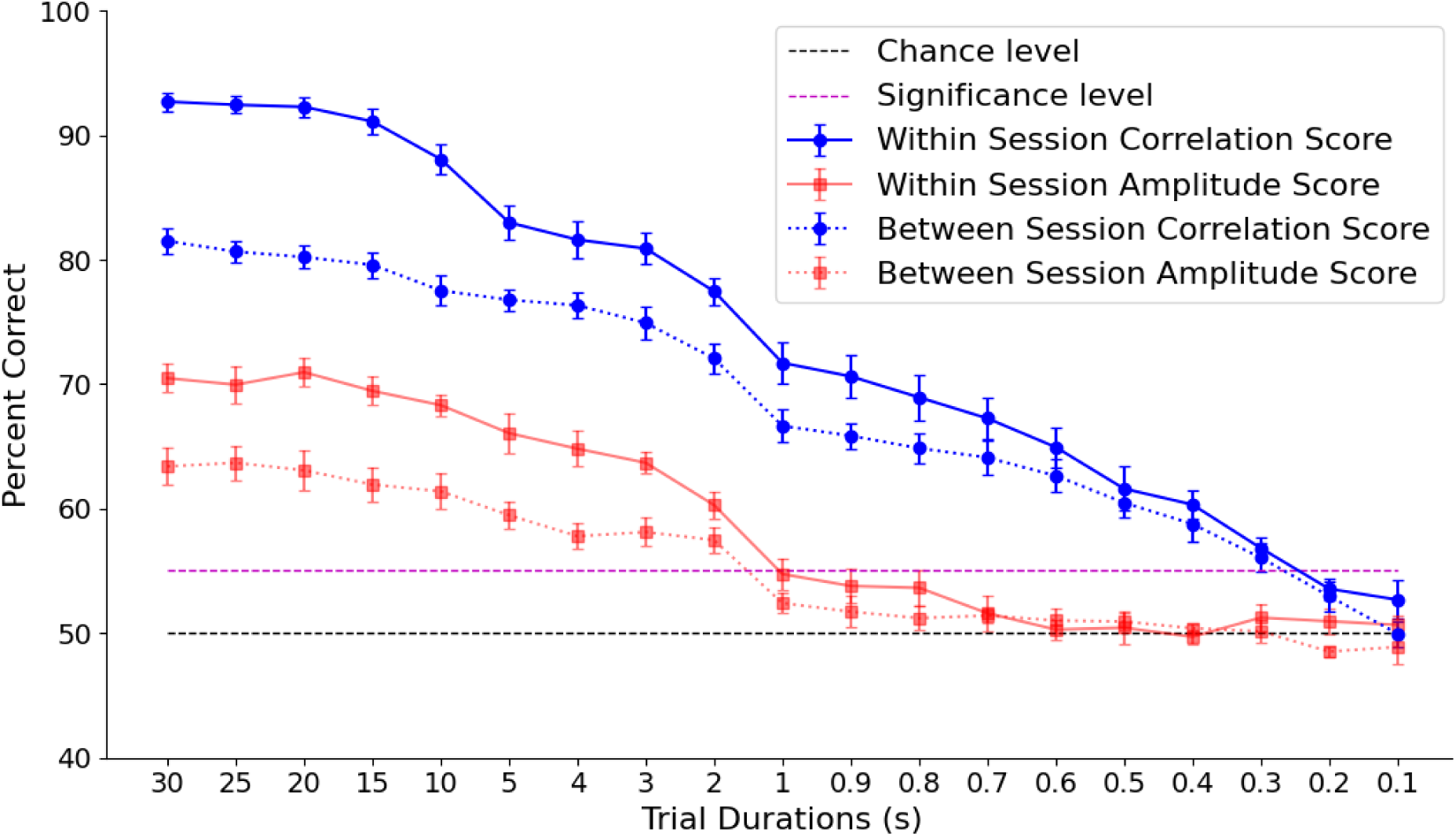
Mean decoding accuracy across electrodes in percent of visual scenes correctly identified as a function of the amount of data used to create the SSVEPs. “Trial duration” refers to the time used to create both the SSVEPs from the scene being tested and the SSVEPs from the scene it is being compared to. Decoding accuracy for correlation and amplitude decoding are shown for within and between sessions decoding. Black dashed line indicates 50% chance level for pairwise decoding and red dashed line indicates the group average percentage correct corresponding to a Bonferroni corrected p-value of 0.00625 for reference. Error bars indicate +/- standard error. The waveform shape contains significantly more information about the visual scene than the amplitude.

#### Determining the amount of flicker time needed to successfully decode

As expected, the average decoding accuracy became worse as less data was used; the fewer segments that are used to create the SSVEP, the lower the signal-to-noise ratio and the more likely it is that the waveform will not have a high enough correlation with the corresponding SSVEP from the second session (or testing day) to distinguish it from other locations. Figure 5 shows the deterioration of decoding accuracy with the shorter durations tested. The Peak-to-peak decoding was significantly worse than the correlation decoding and fell to below significance level with 1 second of data. The decoding using correlation of the waveform shape was consistently better and did not fall to below statistical significance level until only 0.3 seconds of data was used, which corresponds to 3 individual flickers at 10 Hz.

#### Bandpass filtering

For the waveform shape correlation decoding, the band-pass filtered data was always considerably worse than decoding using the raw unfiltered data. This was true for both the within session and between session decoding (figure 6). This indicates that information about the visual scene is embedded in multiple harmonics of the 10 Hz flicker frequency (i.e. 10, 20, 30, 40Hz …. up to over 100 Hz), and multiple harmonics contribute to the highly robust decoding described above.

**Figure 6:**
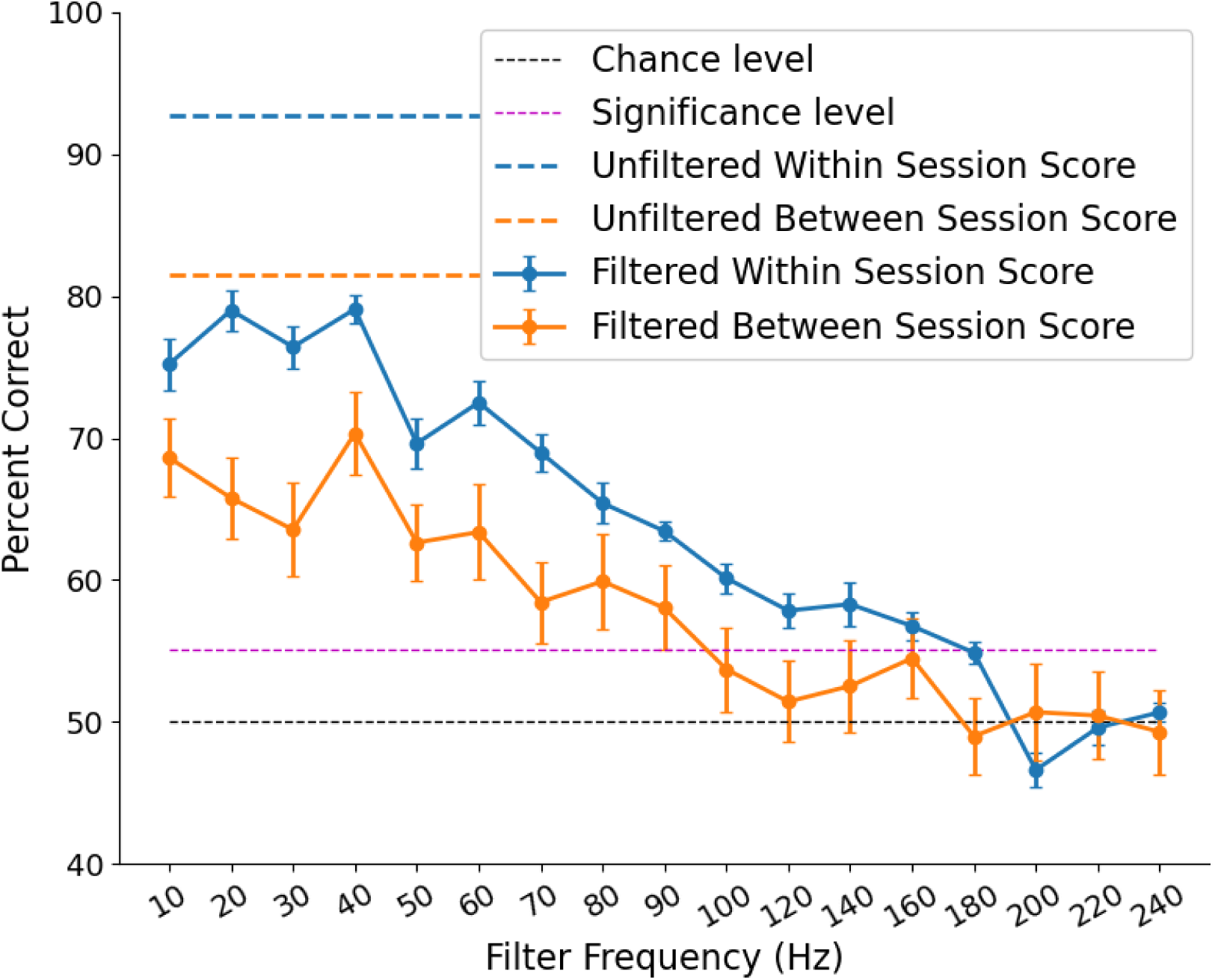
Mean correlation decoding accuracy across electrodes in percent of visual scenes correctly identified after band-pass filtering the data at the various harmonics of the 10 Hz flicker frequency. Decoding accuracy is shown for within and between sessions decoding. Blue and orange dashed lines indicate decoding accuracy from the unfiltered data for within and between sessions respectively. Black dashed line indicates 50% chance level for pairwise decoding and red dashed line indicates the group average percentage correct corresponding to a Bonferroni corrected p-value of 0.00625 for reference. Error bars indicate +/- standard error. For both within and between sessions the filter frequency with the highest decoding accuracy is 40 Hz.

Interestingly, the highest decoding accuracy for any single filter frequency was 40 Hz for both within (79.98%) and between sessions (70.44%). This indicates that resonant gamma band oscillations were evoked (or entrained) as a harmonic of the 10 Hz flicker and contained more information about the visual scene than activity at the fundamental frequency (10 Hz).

With progressively higher frequencies of band-pass filter the decoding accuracy deteriorated until reaching chance level for frequencies above approximately 180 Hz; above this frequency the signal no longer contained any information about the visual scene.

#### Comparison of decoding using endogenous alpha power without flicker

The mean decoding scores using peak endogenous alpha power for each electrode were O1 = 53.25%, O2 = 58.58%, P3 = 55.08%, P4 = 55%, P7 = 54.75%, P8 = 53%.

Only electrode O2 was slightly above the 55.1% accuracy needed to achieve significance at the Bonferroni corrected value (*p* < 0.00625). Therefore, it was not possible to reliably decode the visual scene using the amplitude of endogenous alpha oscillations at the single electrode level with the data from the 4 second baseline period. Electrode O2 was slightly above chance level, indicating that endogenous alpha power is a weak source of information about the visual scene. Visual inspection of the average FFT from the baseline period for each location is informative as to why this is the case (figure 7). Most visual scenes result in a very similar alpha peak, except “Near Space 1”, which was (on average) noticeably higher than the others. “Near space 1” was a blank white wall, viewed from approximately 1 metre. It is unsurprising that a featureless visual input would result in significantly larger alpha oscillations than the other more interesting scenes, and this is most likely the source of the limited decoding accuracy, i.e. the algorithm was able to distinguish the white wall from the other scenes simply because this is the only scene which results in higher alpha power than the others (in some participants).

**Figure 7:**
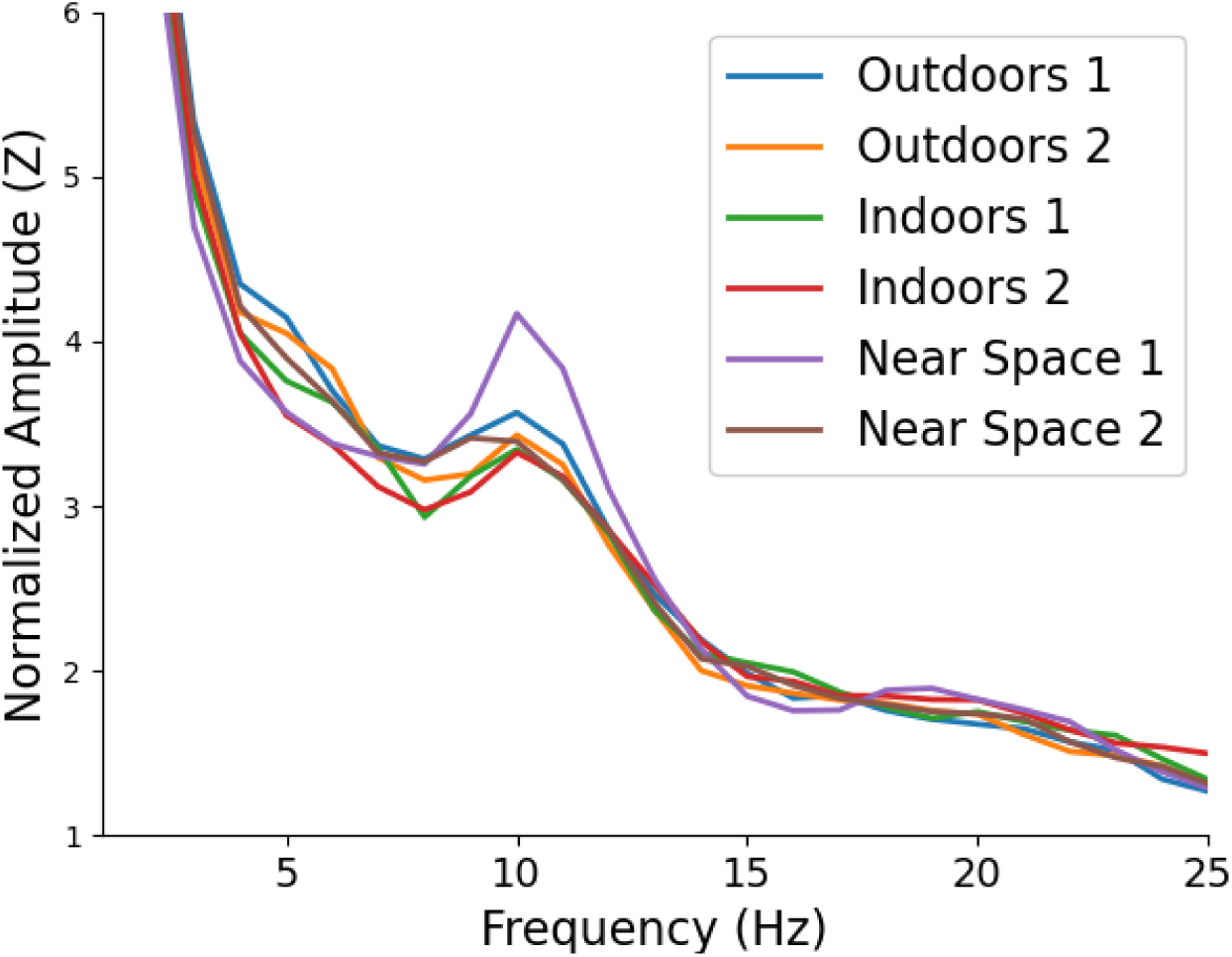
Group average FFT spectra from the 4 seconds of data before the flicker began for each of the 6 visual scenes. All visual scenes resulted in a similar alpha peak, except “Near Space 1”, which was a blank white wall viewed from approximately 1 metre away.

#### Relationship between individual alpha frequency and decoding accuracy

The individual alpha frequency taken from the eyes-closed basement measurement at the beginning of the testing session ranged between 8.25 and 12.5 Hz (average = 10.275 Hz, standard deviation = 0.843 Hz). This resulted in a maximum of 2.5 Hz difference between the 10 Hz flicker frequency and the IAF of participants, with 5 participants showing an IAF of 10 Hz (within an accuracy of +/- 0.25 Hz). There was no significant correlation between SSVEP waveform shape decoding accuracy and IAF difference from 10 Hz (r = -0.08 session 1, r = - 0.13 session 2, all *p* values >0.05). Likewise, there was no significant correlation between SSVEP amplitude decoding accuracy and IAF difference from 10 Hz (r = 0.14 session 1, r = - 0.15 session 2, all *p* values >0.05).

#### Relationship between decoding accuracy across days and differences in luminance

The luminance measurements showed a wide range of absolute differences between day one and day two, especially for the scenes primarily illuminated by natural light. For the scene “Outdoors 1” the maximum absolute difference in LUX values between sessions was approximately 5000 LUX. The scene “Indoors 1” also showed a high degree of day-to-day variability of up to 4000 LUX, due to the large windows and the majority of the scene being illuminated by natural light. All other scenes showed a maximum variability of 100 - 200 LUX. In all cases there were a wide range of LUX differences between sessions with several participants having recordings on 2 days where there was almost no difference between luminance.

There was no significant correlation between the SSVEP waveform shape decoding accuracy for each participant at each location and the difference in LUX values across sessions (Outdoors 1, r = -0.11; Outdoors 2, r = -0.07; Indoors 1, r = -0.2; Indoors 2, r = 0.02; Near Space 1, r = 0.19; Near Space 2, r = 0,09, all *p* values >0.05). Likewise, there was no significant correlation between the SSVEP amplitude decoding accuracy and the difference in LUX values across sessions (Outdoors 1, r = 0.3; Outdoors 2, r = 0.22; Indoors 1, r = 0.19; Indoors 2, r = -0.05; Near Space 1, r = 0.26; Near Space 2, r =-0.15, all *p* values >0.05). The lack of any significant correlation indicates that the lower decoding accuracy between sessions (relative to within sessions), and the variability between participants, is not primarily due to differences in luminance, and therefore the decoding is robust against differences in luminance.

#### Decoding individual participant identity

When applying the same decoding procedure to identify each participant with their unique SSVEP waveform shape (within session), the results were highly successful. Looking at each of the 20 participants individually, the median decoding accuracy across locations and electrodes was always above 89% (median: 97.8%, min: 89.47%, max: 100%), and the most common (modal) score for any one electrode or location, was 100%, with 77.7% of electrode/location pairs giving perfect decoding of the individual out of the 20 possible options. All EEG electrodes gave very similar median decoding accuracies (O1, 98%; O2, 98.7%; P3, 99.1%; P4, 99.1%; P7, 97.8%; P8, 98%). This demonstrates that for any one visual scene, the SSVEP of each individual is both unique and stable across time (in this case within one testing session). Figure 8 shows SSVEPs from 20 participants for one visual scene for two sessions.

**Figure 8:**
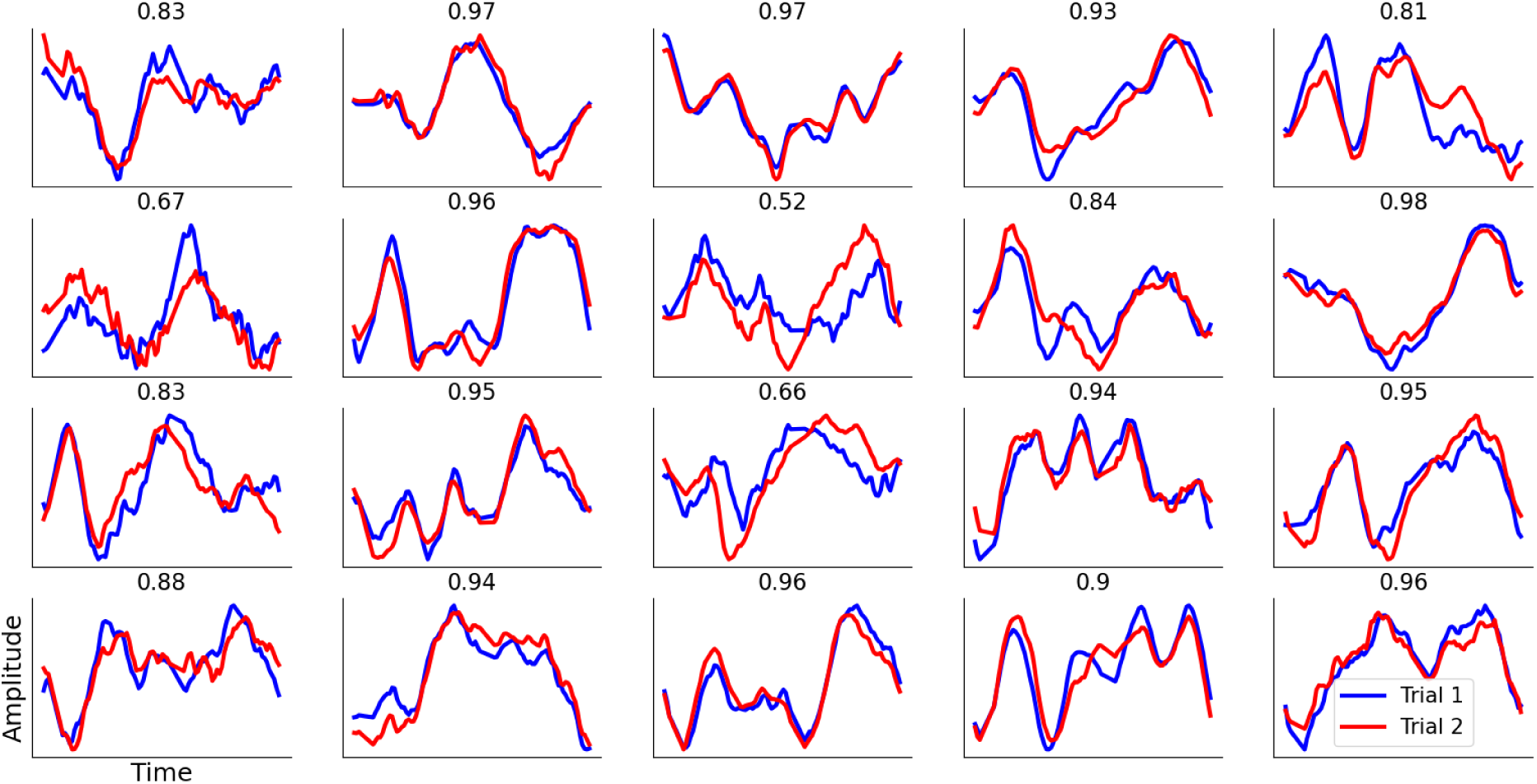
SSVEPs from all 20 participants in experiment 1 from one visual scene (electrode O1). Blue lines are the first trial; red lines are the second trial recorded at a later time on the same day. Most SSVEPs have highly similar waveform shapes; the correlation between the two (Pearson’s r) is shown above each plot. Each participant has a unique SSVEP waveform shape despite all participants standing in the same location and fixating on the same point in the distance.

When decoding the participant across separate days the decoding accuracy fell, but the most common (modal) decoding score across all electrode/location pairs was still 100%. Looking at each participant separately, the median decoding accuracy across locations and electrodes was always above 60% (median: 90.1%, min: 60.9%, max: 97.8%), with 49% of electrode/location pairs giving perfect (100%) decoding.

### Experiment 2

#### Comparison of Decoding accuracy across flicker frequencies

At the group level, using the correlation-based decoding algorithm and averaging across the 6 EEG electrodes, we obtain mean accuracies of 86.4 % for 1 Hz flicker, 87.7% for 10 Hz flicker and 82.5 % for 40 Hz flicker (figure 9). These were all considerably higher than the 54.17% accuracy needed to achieve significance at the Bonferroni corrected value (p = 0.0166). Specifically, Z scores were 19.01, 19.7, 17.01, and *p* values were 6.28 × 10⁻⁸¹,1 × 10⁻⁸⁶, 3.38 ×10⁻⁶⁵, for 1, 10 and 40 Hz respectively. 3 participants got perfect decoding scores across all electrodes and locations for both 1 and 10 Hz (although a lower score than experiment 1 was expected as this was only 10 seconds of data). The mean score for the 10 Hz condition in experiment 2 (87.7%) was very similar to the first 10 seconds of data in experiment 1 (88.06 %), replicating the effect in a second group of participants. While 10 Hz flicker gave the highest mean decoding accuracy, the permutation tests showed that this was not significantly better than 1 Hz (Z = 0.37, *p* = 0.356) or 40 Hz (Z = 1.59, *p* = 0.06)

**Figure 9:**
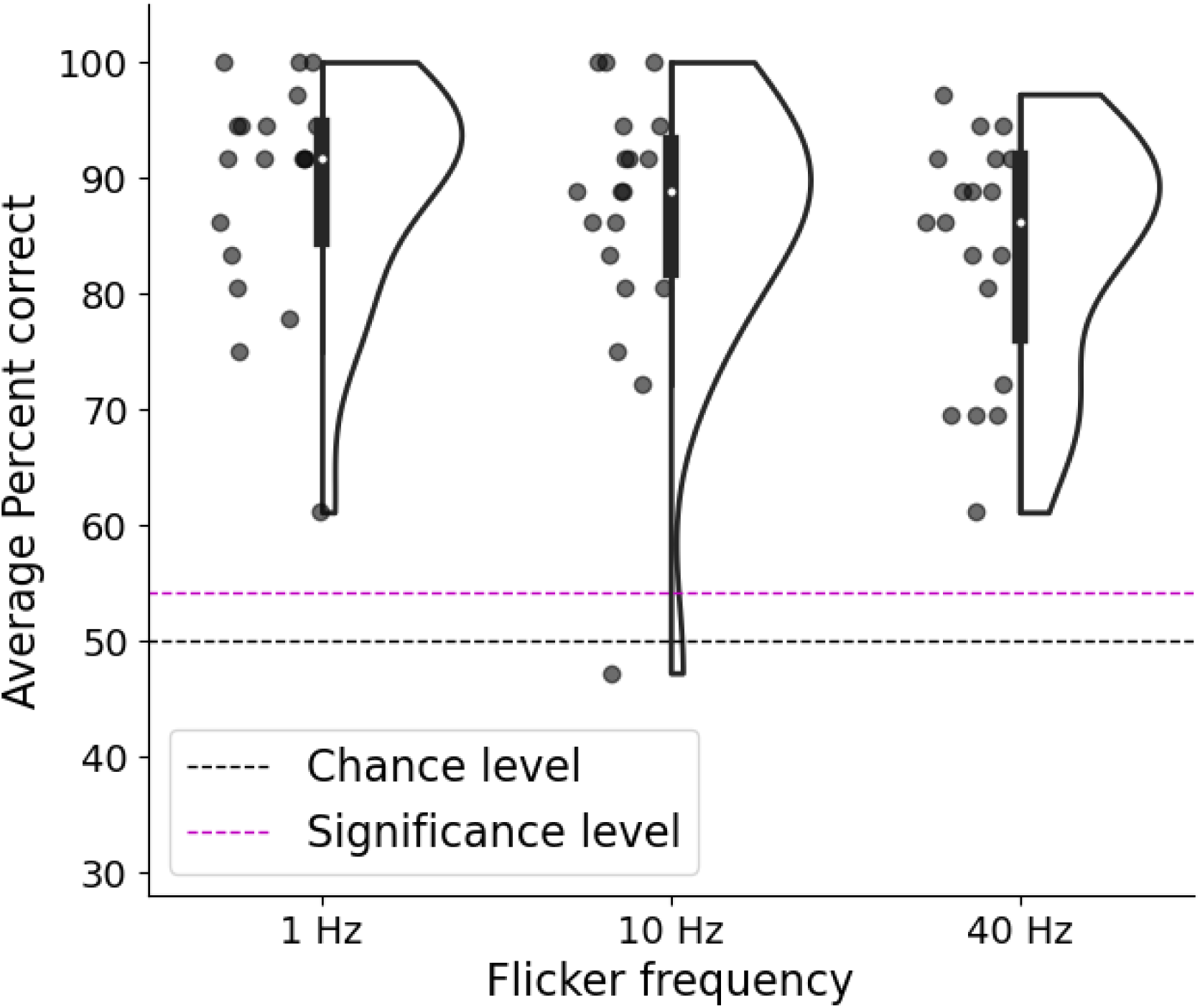
Mean correlation decoding accuracy across electrodes in percent of visual scenes correctly identified for 1, 10 and 40 Hz visual flicker. Each dot indicates one participant, white dot indicates median score, black dashed line indicates 50% chance level for pairwise decoding and red dashed line indicates the group average percentage correct corresponding to a Bonferroni corrected p-value of 0.0166.

#### Band pass filtering

For all 3 flicker frequencies, all band pass filtering resulted in a lower decoding accuracy than using the unfiltered data (figure 10). In all cases, as the filter frequency increases, the decoding accuracy eventually falls to chance level when the corresponding frequency components of the SSVEPs no longer contain any information about the visual scene.

**Figure 10:**
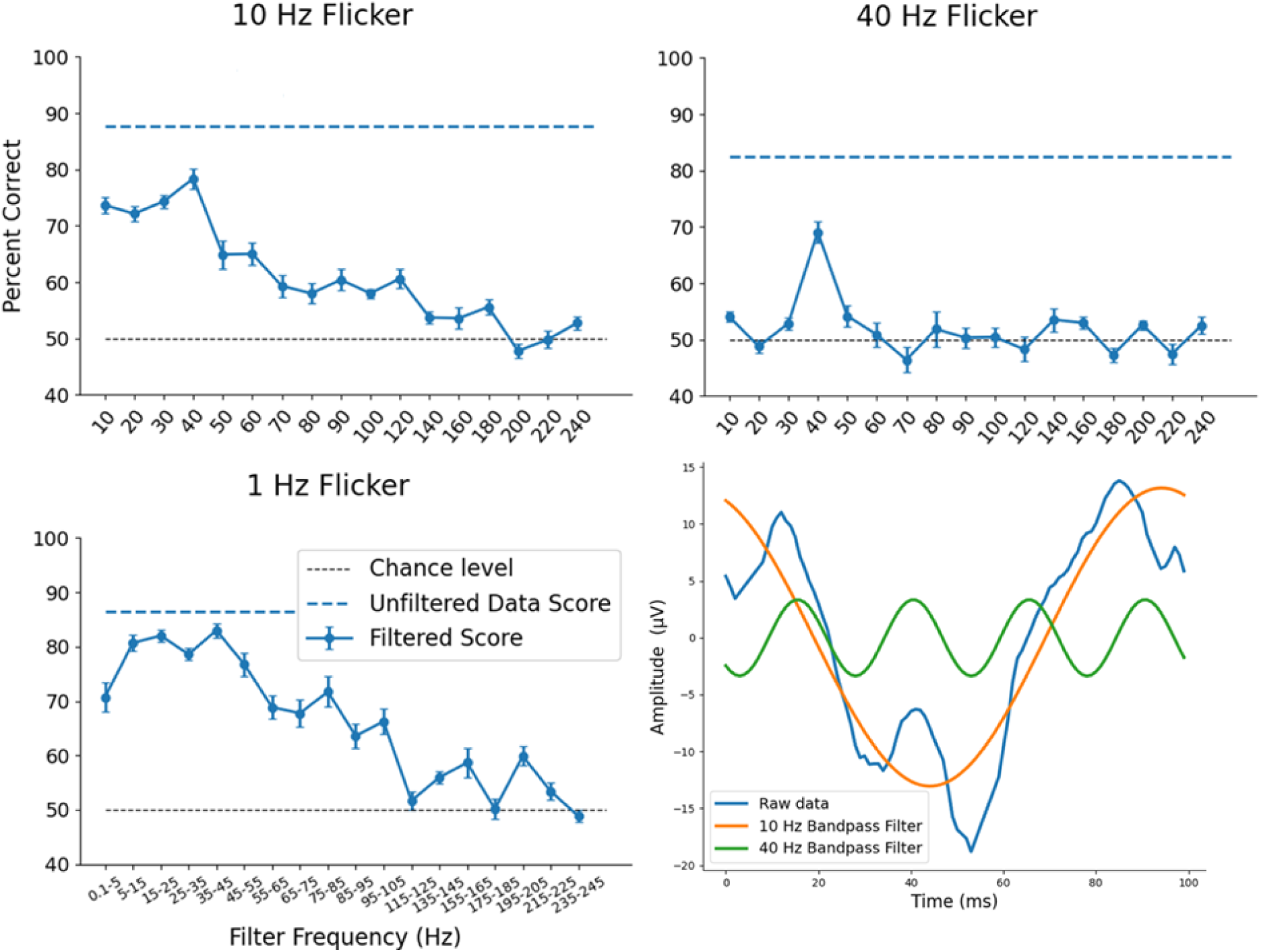
Mean correlation decoding accuracy across electrodes from experiment 2, in percent of visual scenes correctly identified after band-pass filtering the data, as a function of band pass filter centre frequency. Results are shown for 10, 40 and 1 Hz visual flicker. Wider band pass filters were used for 1 Hz flicker as a wider range of frequencies were evoked. Blue dashed lines indicate decoding accuracy from the unfiltered data. Black dashed line indicates 50% chance level for pairwise decoding. Error bars indicate +/- standard error. In all cases the unfiltered data gave the highest decoding accuracy, and band pass filters centred around 40 Hz consistently gave the highest decoding accuracy after filtering. Bottom right; example 10 Hz SSVEP from one participant showing the unfiltered data, and SSVEPs created after 10 Hz and 40 Hz band pass filtering.

For the 10 Hz flicker, the highest decoding accuracy for any of the band-pass filter frequencies was 40 Hz, replicating the results of experiment 1. For the 40 Hz flicker, the best decoding accuracy after band-pass filtering was at the fundamental (40 Hz), with all other band-pass filters resulting in decoding accuracies close to chance level. This indicates that the SSVEPs resulting from 40 Hz flicker are generally close in shape to a sine wave, meaning the signal is only moderately affected by the 40 Hz band-pass filter, whereas the signal is largely removed by all other band-pass filters. Visual inspection confirmed this to be the case.

For the 1 Hz flicker, a wider band-pass filter of 10 Hz was used, centred around multiples of 10 Hz. Decoding accuracies were fairly high across 5-55 Hz, indicating that a range of evoked oscillations contain information about the visual scene across the alpha, beta and gamma bands, with the best decoding accuracy being from the band-pass filter centred around 40 Hz.

## Discussion

We demonstrate that SSVEPs generated from LCD glasses can be used to decode real-world visual scenes with high accuracy. Although a wide range of harmonics contain information about the visual scene, the gamma band is of particular importance. Decoding was successful largely due to the unique waveform shape (rather than the amplitude). The unique SSVEP was reproducible within individuals across time - even across sessions on different days. Decoding was robust: accuracy was above chance with only 300 ms of data, and with enough data performed perfectly (at 100%) in many participants; decoding was possible with single electrodes, indeed with any electrode we tested; participants were standing freely without needing to sit in a confined lab; decoding was resilient to eye blinks and small movements; decoding worked well across sessions within a day and across sessions on different days; decoding was resilient to changes in the visual scene such as luminance and decoding accuracy was similar when comparing different groups of participants (10 Hz blocks in Experiments 1 and 2). This robustness indicates a central role for oscillatory activity - particularly within the gamma band - in encoding complex visual information under naturalistic, real-world conditions.

### SSVEP waveform shape differs strongly across scenes and participants

We used flicker to drive the visual system at specific frequencies, to extract more information than by simply passively measuring neural oscillations. Highly distinct waveforms were evoked for each scene and individual, likely reflecting an idiosyncratic combination of cortical generators, each producing an electrical dipole time locked to the flicker, and each having distinct orientations, phases and time courses, summing together at the scalp level. An alternative (and not mutually exclusive) explanation is that the stimuli are being processed differently between people due to different experiences or associations with the stimulus set. Although SSVEPs are sometimes considered to be reflect sensory processing in early visual cortex (Regan, 1989). SSVEPs can be shaped by experience (e.g. McTeague et al., 2015). In any case, the observed inter-individual variability means that a traditional group-level average across participants, as commonly used in ERP studies, would reduce the information in the signal.

### Reliable decoding with minimal data

By reducing the amount of data used for both the “training” and “test” SSVEPs we found that accurate scene classification was possible using as little as 0.3 seconds of data (i.e. 3 individual flickers at 10 Hz for “training” and 3 individual flickers for the “test” data), at occipital electrodes. This is a remarkably high amount of information per unit time compared to traditional neuroimaging methods such as resting-state EEG without flicker, conventional ERPs and fMRI.

### Robustness to artefacts and movements

The SSVEP decoding was highly robust against eye-blinks and small head movements. So many segments are averaged, and corrupted segments were so rare, that there was no meaningful difference in accuracy with and without rejection of segments with blinks or movement artefacts. Any artefacts or environmental noise not time locked to the flicker averages to zero given enough segments, meaning a reliable and consistent SSVEP from a single electrode can be collected even in mobile participants. This finding extends prior work showing that clean SSVEPs can be acquired during walking (Dowsett et al., 2020).

### Topography of decoding

Decoding accuracy was highest at occipital electrodes, where visual SSVEP amplitudes were largest and the signal-to-noise ratio is high. Nonetheless, accuracy was high across all electrodes. The widespread ability to decode - even from the EOG electrodes - may be due to volume conductance of the neural signal combined with the high signal-to-noise ratio of SSVEPs. It may also be the case that signals from the retina and optic nerve contribute to the EOG. Future work will be required to disentangle the relative contributions of cortical and pre-cortical sources to decoding accuracy at these sites.

### Waveform shape decoding out-performed decoding from amplitude or resting alpha oscillations

SSVEPs have been a standard tool in cognitive neuroscience (Norcia et al., 2015) since they were first introduced over 90 years ago (Adrian & Matthews, 1934). Conventional SSVEP analyses have typically used a frequency transform and have then focused almost exclusively on amplitude at the flicker frequency or a combination of harmonics as the primary output measure. Recent research has emphasised the importance of waveform shape when interpreting neural oscillations (Cole & Voytek, 2017). Here we demonstrate that the waveform shape of an SSVEP (which is independent of amplitude) is a rich source of information about the visual scene.

### Effects of flicker frequency

Normal mental function relies upon recurrent activity dynamically adapting across many neural assemblies. Different frequency bands of oscillations may become particularly prominent in the signal depending on the species investigated (mouse/ human), data acquisition method (EEG/MEG/single cell), stimulus presentation (varying in size/ motion/ ecological validity) and cognitive state. Here we found that the frequency information encoded in the EEG in response to natural scenes depended on the flicker frequency, but not with a one-to-one mapping. Experiment 2 reproduced the effect seen in experiment 1 but with only 3 separate visual scenes, and compared the relative decoding accuracy of 1, 10 and 40 Hz flicker. We achieved significantly higher than chance decoding with all frequencies, with no significant differences between them.

### Broad-band signals in the SSVEP

The unique waveform shape of each SSVEP can be decomposed into various frequencies. To investigate which frequencies contribute to the decoding accuracy, we band-pass filtered the data at the various harmonics and ran the decoding algorithm again for each one. The most striking finding was that the decoding accuracy was highest for the broadband unfiltered data in all cases. Therefore, by filtering SSVEP data, or only focusing on one harmonic of an FFT, a significant amount of information in the visual signal is being lost. This is an advantage of the current time-domain waveform-shape analysis compared to the conventional frequency-domain.

### Highest decoding accuracy within the 40 Hz band

Band-pass filtering data before creating SSVEPs showed that decoding accuracy peaked consistently in the gamma band, specifically at 40 Hz. This was replicated across 5 separate data sets: the within and between sessions in experiment 1, and the 1, 10 and 40 Hz flicker in experiment 2. This implies a significant role of gamma band activity in the neural representation of visual scenes. This could be either because the visual flicker is driving neural circuits which have a resonant time course of approximately 25 ms, or because ongoing gamma band neural oscillations are being entrained to the visual input, or a combination of the two.

During 40 Hz flicker, it was particularly prominent that the band-pass filter with the greatest amount of information was 40 Hz itself, indicating that most of the information is encoded in the phase of an approximately sinusoidal wave. This may either be due to a phase shift of the neural activity itself, or a different combination of cortical sources resulting in a phase shift at the scalp level.

Although decoding was most successful in the gamma band, the largest SSVEP after band-pass filtering was generally in the alpha band. Alpha-band oscillations are often implicated in visuospatial processing but may critically support processes that were not taxed in the current experiment or may not be optimal for decoding natural scenes: oscillations in the alpha band may reflect multiple dissociable mechanisms (Bonnefond & Jensen, 2025; Cruz et al., 2025), for example not predicting discrimination sensitivity but rather subjective reports (Benwell et al., 2022) or attentional gating (Peylo et al., 2021).

### Oscillations underlying natural scene processing

Gamma oscillations in visual cortex have been implicated in the free viewing of natural scenes, although it is controversial whether the gamma band is the dominant frequency in naturalistic scenes (Hermes et al., 2015a). It has been suggested that gamma band rhythms not visible in the power spectra may nonetheless have a function in natural vision (Brunet et al., 2014); the gamma range component of SSVEPs may be a useful metric in this regard as the high signal-to-noise ratio allows even very weak phase-locked gamma band responses to be observed that would otherwise be obscured by the broadband signal.

Alpha and gamma oscillations may play complementary roles. Local field potential recordings of naturalistic scenes from anaesthetised monkey V1 suggest information streams may be encoded in parallel (multiplexing) across alpha and gamma bands (Mazzoni et al., 2011), which may function to keep feedforward and predictive feedback signals separate (van Kerkoerle et al., 2014). More recent human EEG work found that incoherent videos (although presented in small apertures, unlike the full-field stimuli in the current study) were encoded best in the gamma band whereas coherent videos were decoded best in the alpha band (Chen et al., 2023, 2025).

There may be key differences between looking at responses in the brain at rest, as compared to when driven by flicker. The flicker response indicates the brain’s excitability to light (i.e. photic) stimulation at different frequencies and how that excitability manifests in resulting oscillations. Further work is necessary to fully uncover when the response to flicker reveals aspects of brain processing that are relevant for cognition.

### No relationship between individual alpha frequency and decoding accuracy

We did not find any relationship between how close the flicker frequency was to the participant’s individual alpha frequency (IAF) and decoding accuracy, contrary to our hypothesis. Previous research has shown visual flicker at individual alpha frequency drives greater connectivity across regions (Jaeger et al., 2023) and has distinct effects on various sources of cortical alpha (Nuttall et al., 2022). However, these studies actually administered visual flicker at the individual alpha frequency (to the nearest 0.25 Hz), rather than always flickering at 10 Hz and measuring the distance to IAF, as we did here, and this may explain the lack of effect.

### No relationship between day-to-day differences in luminance and decoding accuracy

There was no significant relationship between luminance and decoding accuracy, again contrary to our hypothesis, despite significant differences in luminance across days for the visual scenes of outdoor environments. Differences in decoding accuracy across days may be due to factors such as differences in the EEG signal across sessions and the mental state of the participant (e.g. tiredness, mind wandering).

### Eye movements

A key concern in any EEG experiment is to rule out any effects being due to the execution or preparation of eye movements, which cause artefacts in the EEG signal. It is important to note that in this experiment the participants were passively viewing the stimuli. Recent studies have shown that although when performing attentional and memory tasks, microsaccades were systematically oriented towards salient stimuli, potentially affecting the EEG, strikingly this did not occur during passive viewing conditions (Cruz et al., 2025; de Vries & van Ede, 2025), and so cannot be attributed to the current study where participants were only viewing passively. Similarly, here we did not find any evidence that eye movements were entraining to the flicker. Decoding accuracy was in fact weaker around the EOG electrodes where the muscle artefact is strongest.

### Assessing our decoding in the context of other decoding studies and methods

Previous studies using multivariate decoding of visual ERPs have also recently demonstrated comparable accuracy (albeit with carefully controlled artificial stimuli) showing decoding of stimulus properties like colour, orientation, identity etc. (e.g. Bae & Luck, 2019), and SSVEPs are decoded in some brain-computer interface applications (Saha et al., 2021; Wang et al., 2024). The novel contribution of the current results goes beyond the high accuracy, rather relating to the capability to decode under highly naturalistic conditions, the specific implication of the gamma band in decoding, the discovery of how informative SSVEP waveform shape is, the empirical efficiency (a single electrode with less than a second’s data), and robustness across re-testing.

### Practical significance

Indeed, the manner of our decoding bears substantial practical applications. The slow temporal resolution and slow acquisition rate of fMRI mean even with the latest breakthroughs, fMRI decoding of visual scenes using deep learning necessitates scanning time of at least 2 hours but has usually required at least 20 (Ferrante et al., 2024) leaving some to eschew these for other machine learning techniques (Trammel et al., 2023). The rapid data acquisition time of the flicker glasses technique may be valuable in the future for analysis of large populations/datasets. Additionally, we note that compared to deep learning and fMRI the current setup costs less money, which is an important factor in global initiatives hoping to introduce cutting-edge neuroscience to developing countries, increasing diversity of research and researchers (Cirulli et al., 2025; Shen et al., 2021). The potential of decoding with short periods of data collection in mobile environments makes it relevant for clinical and even domestic use, and the use of low numbers of electrodes and decoding across all positions tested is relevant for the design of wearable systems using e.g. ear electrodes (Debener et al., 2015).

### Limitations and Outstanding questions

In the current study we presented rhythmic flicker at specific frequencies. Several studies have demonstrated distinct neural activity resulting from rhythmic and arrhythmic visual flicker in both EEG (Notbohm et al., 2016) and fMRI (Parkes et al., 2004). The assumption is that rhythmic stimulation is more likely to entrain ongoing neural oscillations, but the effect could also be due to neural circuits resonating as a result of rhythmic stimulation without any endogenous oscillations being entrained. Future studies should test if the decoding is as effective if the visual flicker is arrhythmic, and if not, the extent to which the rhythmicity of the flicker affects the decoding accuracy.

A related unknown factor is the exact combination of low-level visual properties which were being used for decoding. We have shown that luminance is not a significant factor, but many other factors such as depth, colour, edges, angles etc. may be relevant. Ultimately, we would aim to replicate the results of this experiment with a much larger number of visual environments and begin to correlate low level visual properties with various elements in the SSVEP waveform.

Furthermore, there would be a benefit to recording 64 channels with the current paradigm (as well as with MEG) to enable source reconstruction of the signal. Future studies should pursue both lab-based studies with large numbers of electrodes (and multiple imaging modalities), as well as the minimalist real-world studies, to maximize the benefit of this method.

